# Regional Collapsing of Rare Variation Implicates Specific Genic Regions in ALS

**DOI:** 10.1101/375774

**Authors:** Sahar Gelfman, Sarah Dugger, Cristiane de Araujo Martins Moreno, Zhong Ren, Charles J. Wolock, Neil A. Shneider, Hemali Phatnani, Elizabeth T. Cirulli, Brittany N. Lasseigne, Tim Harris, Tom Maniatis, Guy A. Rouleau, Robert H. Brown, Aaron D. Gitler, Richard M. Myers, Slavé Petrovski, Andrew Allen, Matthew B. Harms, David B. Goldstein

## Abstract

Large-scale sequencing efforts in amyotrophic lateral sclerosis (ALS) have implicated novel genes using gene-based collapsing methods. However, pathogenic mutations may be concentrated in specific genic regions. To address this, we developed two collapsing strategies, one focuses rare variation collapsing on homology-based protein domains as the unit for collapsing and another gene-level approach that, unlike standard methods, leverages existing evidence of purifying selection against missense variation on said domains. The application of these two collapsing methods to 3,093 ALS cases and 8,186 controls of European ancestry, and also 3,239 cases and 11,808 controls of diversified populations, pinpoints risk regions of ALS genes including *SOD1, NEK1, TARDBP* and *FUS.* While not clearly implicating novel ALS genes, the new analyses not only pinpoint risk regions in known genes but also highlight candidate genes as well.

## Introduction

Amyotrophic lateral sclerosis (ALS) is an adult-onset neurodegenerative disease characterized by progressive motor-neuron loss leading to paralysis and death, most often from respiratory failure. Roughly 60-70% of familial and 10% of sporadic cases have an identifiable mutation in a known causal ALS gene, the majority of which are repeat expansions in *C9ORF72* and point mutations in *SOD1^1^*. Recent efforts in gene discovery, largely driven by advances in sequencing and identification of rare variants, have implicated and confirmed several new genes in ALS pathogenesis including *TBK1, NEK1, ANXA11* and *CCNF*^2–10^. Despite this progress, the majority of sporadic cases still remain to be resolved genetically.

The now established paradigm for case-control analyses of exome or genome sequencing data of complex diseases and traits involves a gene-based collapsing framework in which all qualifying variants in a gene are treated as equivalent. Genes are associated with the trait when they exhibit a significant excess of qualifying variants occurring anywhere in the gene. This approach has implicated disease genes in a growing number of other complex conditions beyond ALS, including idiopathic pulmonary fibrosis (IPF), myocardial infarction (MI) and Alzheimer’s disease^11–13^.

While clearly effective, the power of this approach is limited by the inclusion of benign variants that reduce statistical power. However, for genes where pathogenic mutations are localized to specific regions, such as functional domains, power can be increased by using these regions as the unit for the collapsing analysis. In ALS-associated genes, there are several examples of genes that show regionally localized pathogenic variation. For example, in *TARDBP*, highly-penetrant ALS variants are concentrated in a glycine-rich domain near the C-terminus^14^. Furthermore, the gene *FUS*, which has a similar structure as *TARDBP*, has pathogenic mutations clustering in two regions: exons 13-15 (encoding an Arg-Gly-Gly-rich domain and the nuclear localization signal) and exons 3, 5-6 (encoding Gln-Gly-Ser-Tyr-rich and Gly-rich domains)^15^.

Recognizing that undiscovered ALS-associated genes might similarly have specific domains where pathogenic variants cluster, we now apply two complementary regional approaches to gene collapsing analyses to identify localized signals of rare variation in a data set of 3,093 ALS cases of European ancestry (2,663 exomes and 430 whole genomes) compared with 8,186 controls of matched ancestry (7,612 control exomes and 574 whole genomes). We further apply these analyses to a set of samples of diversified ancestry origins, consisting of 3,239 cases and 11,808 controls. We compare the regional approaches to the standard gene collapsing analysis and highlight the importance of a regional view specifically for ALS genetics.

## Results

### Collapsing analyses using homology-defined protein domains

The standard approach to gene discovery focuses on the burden of rare variants across an entire gene by comparing the frequency of qualifying variants in cases and controls. The qualifying variants can be defined by various criteria such as function and allele frequency (illustrated in figure 1A).

**Figure 1.**
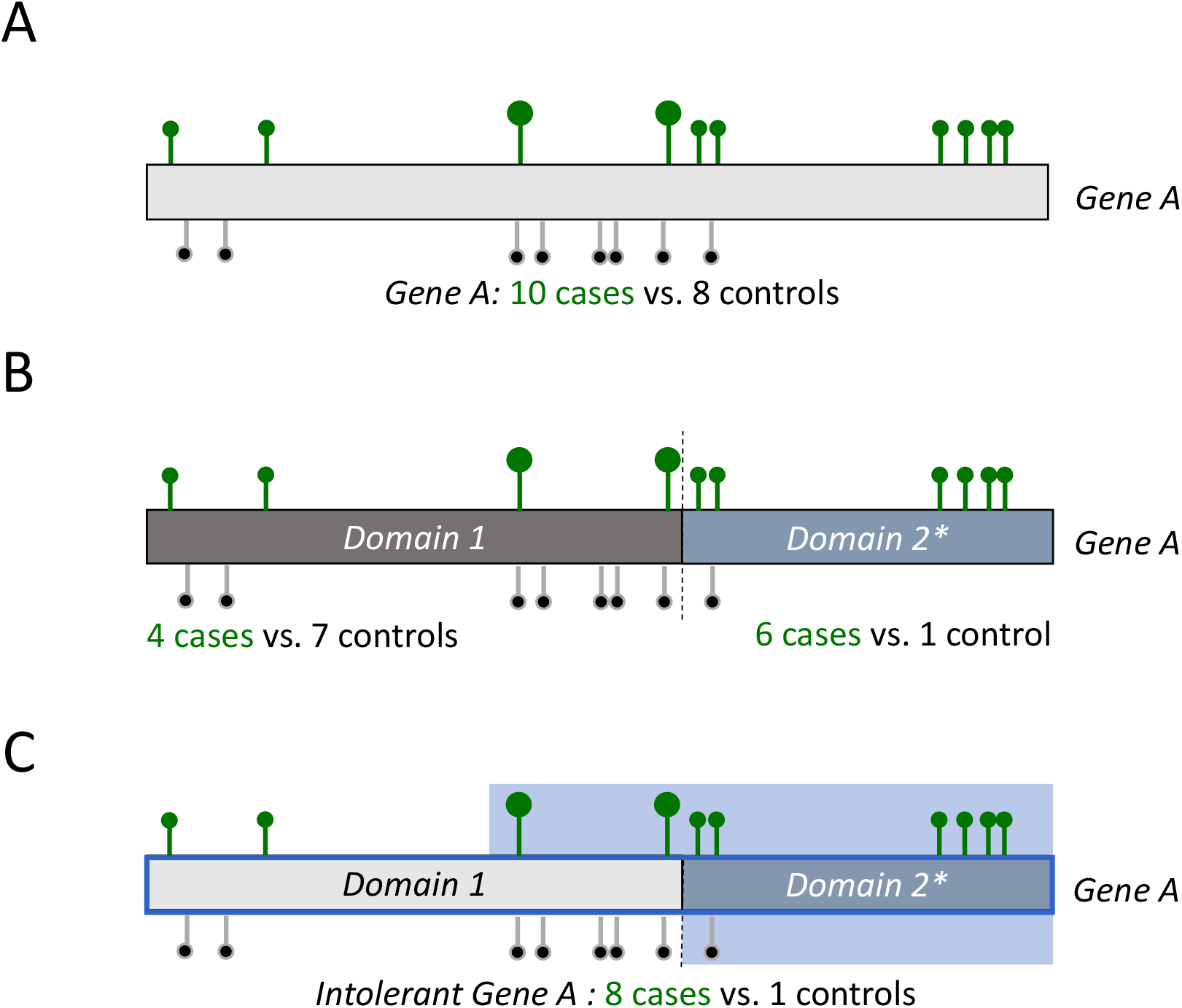
Gene and Regional Collapsing. (**A**) A standard gene-based approach for collapsing analysis of non-synonymous and canonical splice rare variants in cases (green) and controls (black) on example *Gene A.* (**B**) A domain-unit based regional approach where only the domains that are intolerant to functional variation are considered as units for collapsing. (**C**) Intolerance informed gene collapsing: a regional approach to gene collapsing where the unit for collapsing is the entire gene, yet missense variants only qualify for the analysis if they reside in domains that are intolerant to variation (domain 2). Loss-of-function variants (big circles) continue to qualify regardless of whether they reside in a tolerant or intolerant domains of the gene. Bright blue background marks qualifying variants.

In this study, we describe two additional approaches to rare variant collapsing: 1) a regional approach, where the unit for collapsing is not the gene, but rather the functional domains within the gene (figure 1B), and 2) a gene-based approach where the definition of qualifying variants is informed by regional intolerance to missense variation (figure 1C).

**Figure 2.**
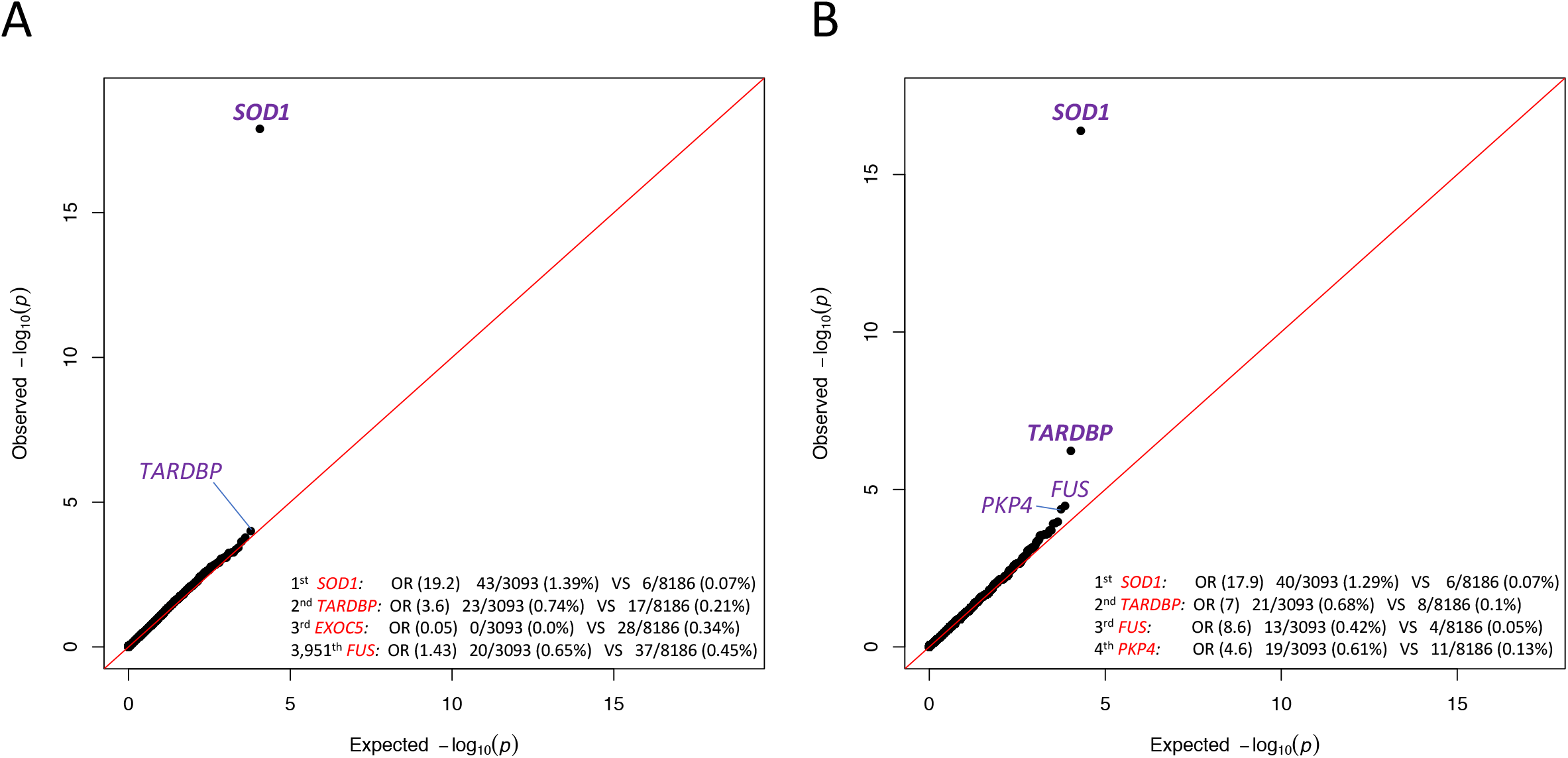
Q-Q plots of gene and domain level collapsing. (**A**) The results for a standard gene level collapsing of 3,093 cases and 8,186 controls. 18,065 covered genes passed QC with more than one case or control carrier for this test. The genes with the top associations and *FUS* gene are labeled. The genomic inflation factor, lambda (λ), is 1.10. (**B**) The results for the domain-based collapsing of 3,093 cases and 8,186 controls. 70,603 covered domains passed QC with more than one case or control carrier for this test. The genes with the top associations are labeled and genome-wide significant genes are in bold. λ=1.046.

We first utilized the standard gene collapsing approach (figure 1A) to identify the burden of rare variants in a set of 3,093 ALS cases and 8,186 controls of European ancestry. The demographic features of our cohort reflect known epidemiological features of ALS, including male predominance and the distributions of age at onset and survival (supplementary Table S1). Qualifying variants were defined as non-synonymous coding or canonical splice variants that have a minor allele frequency (MAF) ≤0.1% in cases and controls (internal MAF) and also a ≤0.1% MAF imposed for each population represented in the ExAC database^16^. High quality control (QC) metrics were further imposed on the variants (see Methods).

Comparing genetic variation across 18,653 protein-coding genes found a genome-wide and study-wide significant (p<4.5 × 10-7) case-enrichment only for *SOD1* (p=1.23×10^−18^, figure 2A), with qualifying variants identified in 43 cases (1.39%) and only 6 controls (0.07%; OR=19.2). *TARDBP* showed the second strongest enrichment (OR=3.6, p=1.02×10^−4^), but with 23 cases (0.74%) and 17 controls (0.21%) it did not achieve genome-wide significance. *FUS* harbored qualifying variants in 20 cases and 37 controls (OR=1.43, p=0.23, figure 2A).

A gene-based analysis evaluating only rare loss of function (LoF) variants was also performed, identifying a genome-wide and study-wide significant case-enrichment of *NEK1* variants (OR=7.35, p=1.85×10^−10^), with 33 cases (1.07%) compared to 12 controls (0.15%, supplementary figure S1). As a negative control, we included a model for rare synonymous variants, and did not observe any genes with significant enrichment. The genomic inflation factor, lambda (λ) for this model was 1.03 (supplementary figure S2).

We hypothesized that genes with clustered mutations that had weak enrichments using this standard gene-based collapsing approach, such as *TARDBP* and *FUS*, could be identified by a collapsing method that uses functional gene regions (i.e. domains) as the unit for collapsing (figure 1B). For this analysis, we utilized a list of 89,522 gene domains covering the human coding sequence, as described previously^17^. In short, the coding sequence of each gene was aligned to a set of conserved protein domains based on the Conserved Domain Database (CDD)^18^. The final domain coordinates for each gene were defined as the regions within the gene that aligned to the CDD and the unaligned regions between each CDD alignment. These domains were then used as the unit for collapsing compared with a standard gene-based collapsing approach (figure 1A and 1B).

This domain-based analysis was performed using the same cohort and coding model as the standard approach (European ancestry, non-synonymous and canonical splice variants, internal and population MAF ≤0.1%). As hypothesized, the top three case-enriched domains reside in ALS genes: *SOD1, TARDBP* and *FUS* (figure 2B). For *SOD1*, a long domain spanning the majority of the coding sequence contains most of the variation found in 1.29% of cases and 0.07% of controls (OR=17.9; p=4.1×10^−17^, figure 2B).

Strikingly, the glycine-rich *TARDBP* domain where known mutations cluster is now identified with genome-wide and study-wide significance (OR=7; p=5.84×10^−7^). Of note, this glycine-rich domain covers exon 6 of *TARDBP* and was not mapped to a conserved domain from the CDD.

The same trend was observed for *FUS*, which shows the third strongest enrichment in this analysis (OR=8.6; p=3.6×10^−5^, figure 2B). Specifically, qualifying variants were identified in 13 cases (0.42%) and 4 controls (0.05%) in the previously reported Arg-Gly rich domain covering exons 13-15, which is also not considered a conserved CDD domain^14^). Although not at genome-wide or study-wide significance, this represents a substantial improvement over the gene-based collapsing approach (OR=1.43, uncorrected p=0.23).

The fourth most case-enriched domain was a conserved armadillo repeat domain spanning exons 12-14 of *PKP4* (plakophilin 4, also known as p0071). Qualifying variants occurred in 0.61% of ALS cases and 0.13% of controls (OR=4.6, p=4.1×10^−5^). While not genome-wide significant, *PKP4* is an interesting candidate gene that has been previously linked to various ALS-related pathways (see Conclusions).

### Gene-wide collapsing analyses informed by regional intolerance to missense variation

As we have demonstrated, domain-based collapsing effectively identifies genes where pathogenic variants are localized to single specific regions (e.g *TARDBP* and *FUS)*, and highlights suggestive candidates for further study (PKP4). However, to identify haploinsufficient genes where truncating variants and sufficiently damaging missense mutations could both contribute to risk of disease, the difficulty lies in determining which missense variants should qualify in the analysis. To address this challenge, we implemented a collapsing approach that leverages regional patterns of intolerance to missense variation (sub-RVIS^17,19^) as a way to prioritize missense variants most likely to result in disease. In this ‘intolerance-informed’ approach, rare missense alleles were counted as qualifying if they resided in gene regions that are intolerant to missense variation, whereas LoF variants were counted as qualifying regardless of location within the gene (figure 1C).

As a measure of intolerance of gene regions, we applied a complementary approach to subRVIS^17^ for when there is limited resolution in the sequence region of interest. This approach uses the observed to expected missense ratio in a domain (OE-ratio), which is equivalent to a domain-based missense tolerance ratio (MTR)^19^. In short, the expected rate leverages the underlying sequence context in the domain, and the observed rate is based on the rate of non-synonymous variants identified in the sub-region of interest based on the ExAC database, release 0.3^16^ (see Methods).

We focus our intolerance-informed gene collapsing approach on domains that have intolerance below the exome-wide 50^th^ percentile (OE-Ratio percentile <50%), thus sub-selecting variants in genic regions that have greater evidence of purifying selection acting against non-synonymous variation. As mentioned earlier, for each gene, variants in these intolerant regions are considered along with LoF variants independent of their location within the gene.

Because intolerant coding regions are expected to have a lower rate of common variation, we included samples from diversified ancestries when applying intolerance-informed gene collapsing. The diversified population approach increased the total number of samples by 3,768, to 3,239 cases and 11,808 controls, thereby increasing the power of the analysis. For this approach, we applied similar rules for qualifying variants, including low population frequency (MAF ≤0.1% imposed for each population represented in ExAC), an internal MAF ≤0.04% (decreased from 0.1% due to a larger control cohort), coding annotation (non-synonymous and splice variants) and high QC metrics, with the additional criteria of residing in the lower 50^th^ percentile of OE-ratio domains.

The genomic inflation factor (λ) of the diversified populations intolerance-informed analysis was 1.14, slightly higher than the European-only cohort used for the standard gene level analyses (figure 2A, λ= 1.1). Yet, this inflation is much lower than for the standard gene based analysis using a diversified population (λ=1.25, supplementary figure S3), demonstrating the advantage of an intolerance-informed approach for reducing the genomic inflation due to variation in tolerant regions.

In this analysis, *SOD1* achieved a slightly better enrichment than in either gene-based or domain-based analyses (OR=20.31; p=4.13×10^−22^, figure 3A).

**Figure 3.**
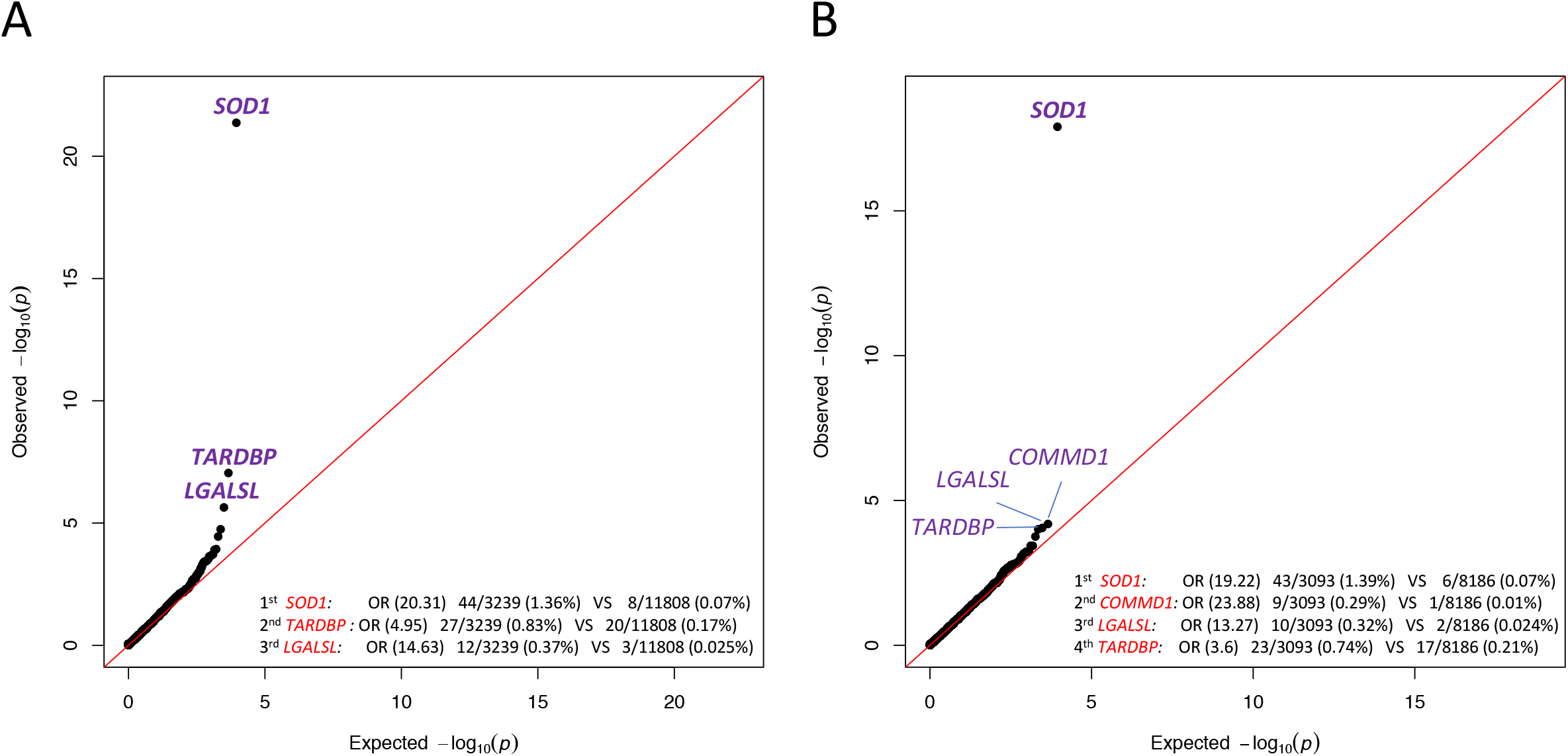
Intolerance informed gene level collapsing with unified/diversified ancestry samples. (**A**) A q-q plot presenting the results of the gene-based intolerance-informed collapsing of 3,239 cases and 11,808 controls from diversified ancestries. Missense variants are aggregated only if they reside in an intolerant domain that is lower than 50^th^ percentile OE-ratio score, while loss-of-function variants are aggregated independent of location. 17,795 genes passed QC with more than one case or control carrier for this test. The genes with the top associations are labeled. λ=1.14. (**B**) A q-q plot of a gene-based intolerance-informed collapsing of 3,093 cases and 8,186 controls of European ancestry. 18,135 genes passed QC with more than one case or control carrier for this test. The genes with the top associations are labeled and genome-wide significant genes are in bold. λ = 1.073.

*TARDBP* also had genome-wide and study-wide significant enrichment (OR=4.95; p=8.77×10^−8^; figure 3A), which presents a considerably-strengthened enrichment signal for *TARDBP* compared to the standard gene collapsing analysis observed in figure 2A (OR=3.6, uncorrected p=1.02×10^−4)^.

*LGALSL* (Galectin-like or Lectin, Galactoside-Binding, Soluble, Like) was the third gene to have a strong enrichment of qualifying variants in cases (OR=14.63; p=2.29×10^−6^; figure 3A) that was not study-wide significant given the models tested. The enrichment of this gene originates from one specific domain that harbors variants for twelve cases (0.37%) and three controls (0.025%) with the addition of an African American and a Latino case over the European-only analysis (figure 2). The target domain harboring all *LGALSL* case-variants is a region comprising 378bp that is mapped to a conserved protein domain intolerant to variation. Notably, *LGALSL* LoF variants were only identified in cases and absent from nearly 12,000 controls. To assess the rate of LoF variants in a larger control population we looked at the ExAC cohort and found three LoF alleles in 60,033 individuals^16^.

### Genome wide associations with age-at-onset

We next examined whether qualifying variants in known ALS genes, or candidate genes identified by our novel approaches, influence age at symptom onset (AAO). We found that *SOD1* variant carriers tended to be younger than the rest of the cohort (52.2 vs. 57.1 years, p=0.059; MW-test). Also, subjects harboring qualifying variants in *ANXA11* showed delayed onset (63.8 years, p=0.037; MW-test), which is consistent with prior studies^9^. No other known ALS genes showed significant influence on AAO.

Interestingly, subjects harboring *LGALSL* qualifying variants showed a mean AAO that is 13 years younger than the rest of the cohort (43.8 years vs. 57.1, p=8.1×10^−4^; Mann-Whitney test). The AAO information was available for 11/12 variant carriers and 2,767/3,239 non-carriers.

The early onset in cases carrying *LGALSL* variants was further validated by a random sampling approach where *LGALSL* carriers’ average AAO was significantly lower than 9,983/10,000 randomly sampled sets of 11 cases (p=0.0017, supplementary Methods).

## Discussion

Here we present a regional approach to rare variant collapsing analyses. This approach has two distinct forms: 1) aggregating rare variants on genic sub-regions defined using conserved protein domain annotations, and 2) aggregating rare variants on a gene unit but using the pattern of purifying selection to identify the most damaging missense variants and combine them with loss of function mutations occurring anywhere in the gene. Both approaches show improved sensitivity for known ALS genes, finding *SOD1, NEK1* and *TARDBP* as genome-wide significant. We also find *FUS’s* Arg-Gly-rich domain within the top three associations in our domain-based regional collapsing, jumping from an insignificant OR=1.43 to a high OR=8.6. These findings underscore the utility of applying a regional approach to ALS genetics, especially in light of similar Gly-rich domains importance in mediating pathologic RNA-protein complexes^20^.

This approach has also implicated a potential new candidate ALS gene, *LGALSL*, encoding the galectin-like protein GRP (galectin-related protein, also known as HSPC159 and lectin galactoside-binding-like protein). We identified a case-enriched intolerant galectin-binding domain (Figure 4A). Interestingly, while the functions of *LGALSL* remain largely unknown, members of the galectin family, including *LGALS1* and *LGALS3*, have been implicated in ALS disease processes and progression. Specifically, *LGALS1* has been identified as a component of sporadic and familial ALS-related neurofilamentous lesions^21^, and is associated with early axonal degeneration in the *SOD1^G93A^* ALS mouse model^22^. Furthermore, homozygous deletion of *LGALSL3* reportedly led to accelerated disease progression and reduced lifespan in *SOD1^G93A^* mice^23^. We also performed an analysis of age at onset that is independent of the predisposition analysis, showing a significant association between carriers of qualifying variants in *LGALSL* and early age at onset. This independent analysis supports *LGALSL* as a candidate ALS gene that might be responsible for a form of ALS with a younger age at onset. Because *LGALSL* carriers were contributed by five different sites and sequenced at two sequencing centers, we cannot exclude hidden variable stratification that might explain the low AAO. Further study of additional *LGALSL* mutation carriers will be required to confirm this observed genotype-phenotype correlation.

**Figure 4.**
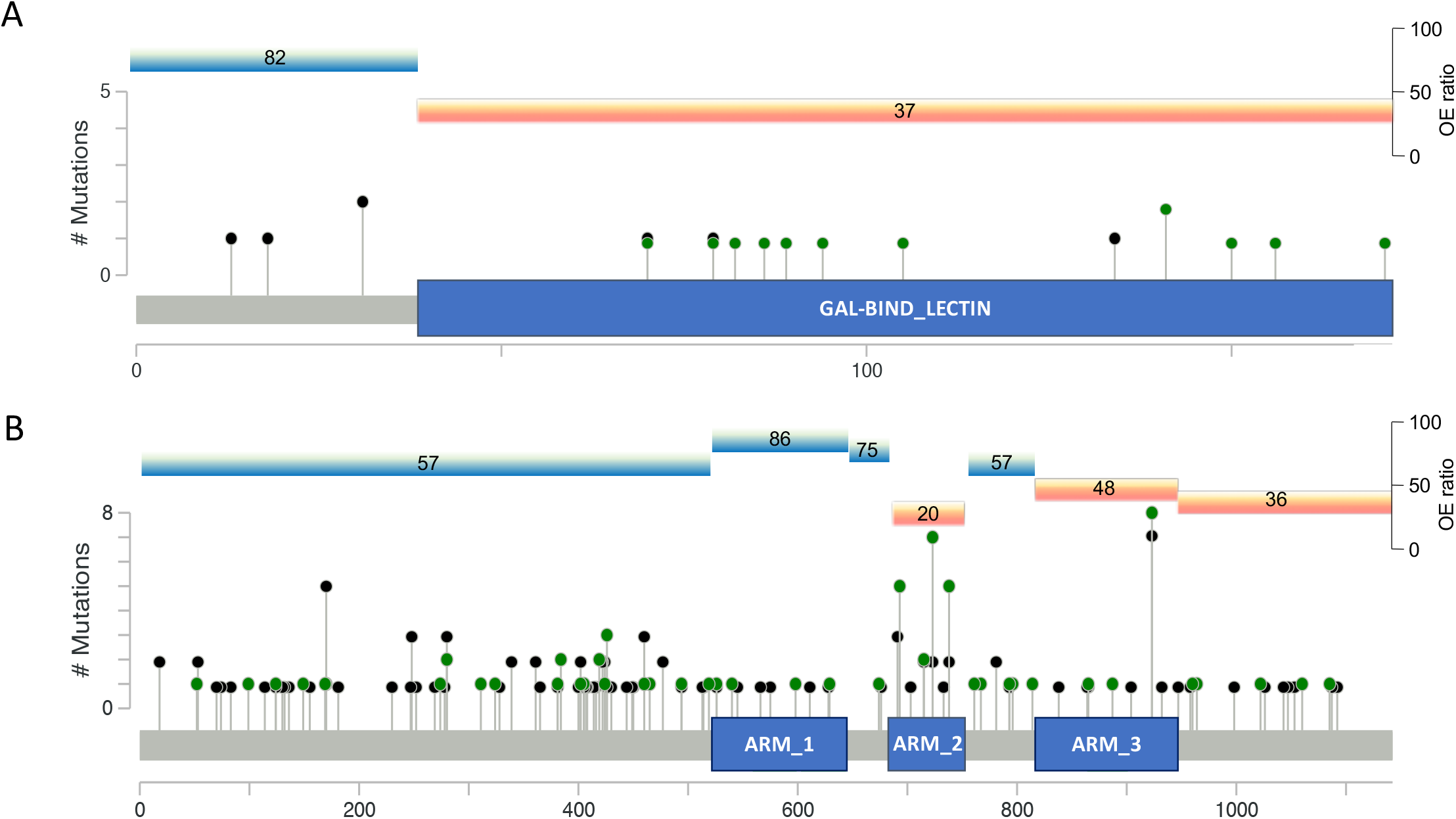
Distribution of functional coding variants across *LGALSL* and *PKP4.* The distribution of *LGALSL* (**A**) and *PKP4* (**B**) coding variants across domains *(LGALSL* transcript ENST00000238875 and *PKP4* transcript ENST00000389757). The y-axis corresponds to the total number of variants identified at a specific location. The blue boxes highlight the (**A**) *LGALSL* carbohydrate binding domain and (**B**) *PKP4* armadillo repeat domain 2 (ARM2) found to be enriched for variants in cases (green) compared to controls (black). Each domain’s OE-ratio percentile is marked above for both tolerant (bright blue) and intolerant (orange) domains.

Regional collapsing analyses also highlighted *PKP4* as a new candidate gene, with a single armadillo repeat domain strongly enriched for qualifying variants in cases (Figure 4B). Evidence supporting *PKP4’s* role in ALS-linked processes including microtubule transport and endosomal processing, in addition to its local translation in ALS-mutant *FUS* granules, all provide evidence in favor of *PKP4* as a risk factor for ALS^24–27^.

This study incorporated both exome and whole genome samples from a large cohort of over 3,000 cases and close to 12,000 controls. Yet, despite these large cohorts, the standard gene collapsing approach identified only *SOD1* and *NEK1* (loss-of-function specific model) as achieving genome-wide significance, and failed to uncover other known signals for ALS risk factors. We were able to capture these signals, along with candidate novel signals, using a regional approach that is informed by missense variation intolerance. That being said, while confirming *TARDBP*, and suggesting

*LGALSL* (0.42% of cases) as a candidate gene, the regional approach was still underpowered with the current sample size to show genome-wide significance for *FUS* and *PKP4* that might reflect true associations. This suggests that even with these signal optimization approaches, larger sequencing studies are required in ALS. We are, however, confident that the continued application of regional approaches to collapsing analyses in ALS and other rare disorders will enable the identification of novel risk factors with small proportions in patient populations, that were previously difficult to identify due to being masked by benign variants in regions that are tolerant to variation.

## Methods

### Subject sources

ALS samples analyzed by whole exome or genome sequencing came from the Genomic Translation for ALS Care (GTAC study), the Columbia University Precision Medicine Initiative for ALS, the New York Genome Consortium, and the ALS Sequencing Consortium (IRB-approved genetic studies from Columbia University Medical Center (including the Coriell NINDS repository), University of Massachusetts at Worchester, Stanford University (including samples from Emory University School of Medicine, the Johns Hopkins University School of Medicine, and the University of California, San Diego), Massachusetts General Hospital Neurogenetics DNA Diagnostic Lab Repository, Duke University, McGill University (including contributions from Saint-Luc and Notre-Dame Hospital of the Centre Hospitalier de l’Université de Montréal (CHUM) (University of Montreal), Gui de Chauliac Hospital of the CHU de Montpellier (Montpellier University), Pitié Salpe□triére Hospital, Fleurimont Hospital of the Centre Hospitalier Universitaire de Sherbrooke (CHUS) (University of Sherbrooke), Enfant-Jèsus Hospital of the Centre hospitalier affilié universitaire de Québec (CHA) (Laval University), Montreal General Hospital, Montreal Neurological Institute and Hospital of the McGill University Health Centre), and Washington University in St. Louis (including contributions from Houston Methodist Hospital, Virginia Mason Medical Center, University of Utah, and Cedars Sinai Medical Center).

### Subject selection criteria

ALS subjects were diagnosed according to El Escorial revised criteria as suspected, possible, probable, or definite ALS by neuromuscular physicians at submitting centers. Subjects were considered sporadic if no first or second-degree relatives had been diagnosed with ALS or died of an ALS-like syndrome. Because screening for known ALS gene mutations prior to sample submission was highly variable across the cohort, gene status was not considered *a priori.* Controls were selected from >45,000 whole exome or genome sequenced individuals housed in the IGM Data Repository. We excluded all individuals with a known diagnosis or family history of neurodegenerative disease, but not all had been specifically screened for ALS.

### Sequencing

Sequencing of DNA was performed at Columbia University, the New-York Genome Center, Duke University, McGill University, Stanford University, HudsonAlpha, and University of Massachusetts, Worcester. Whole exome capture used Agilent All Exon kits (50MB, 65MB and CRE), Nimblegen SeqCap EZ Exome Enrichment kits (V2.0, V3.0, VCRome and MedExome), IDT Exome Enrichment panel and Illumina TruSeq kits. Sequencing occurred on Illumina GAIIx, HiSeq 2000, HiSeq 2500, or HiSeq X sequencers according to standard protocols.

Illumina lane-level fastq files were aligned to the Human Reference Genome (NCBI Build 37) using the Burrows-Wheeler Alignment Tool (BWA)^28^. Picard software (http://picard.sourceforge.net) removed duplicate reads and processed lane-level SAM files to create a sample-level BAM file. Genomes (n=402) from the New York Genome Center were transferred as sample-level BAM files. We used GATK to recalibrate base quality scores, realign around indels, and call variants^29^.

### Samples quality control

The initial sample consisted of 4,149 ALS cases and 15,107 controls. Samples reporting >8% contamination according to VerifyBamID^30^ were excluded. KING^31^ was used to ensure only unrelated (up to third-degree) individuals contributed to the analysis. For controls, where sample collection methods were not known, we excluded samples where X:Y coverage ratios did not match expected sex. For studies where sample collection and processing involved only ALS patients, mismatches were not exclusionary. Further, to be eligible, samples were further subjected to a CCDS 10-fold coverage principal components analysis (PCA) and an ancestry prediction filter (for European ancestry analysis).

### Cohort construction: ancestry prediction

The ancestry classification model was trained using genotyped data from 5,287 individuals of known ancestry and 12,840 well-genotyped and ancestry-informative markers that were limited to the human exome. The model was trained, tested and validated on a set of individuals with ancestry as follows: non-Finnish European(N=2911), Middle Eastern (N=184), Hispanic (N=368), East Asian (N=539), South Asian (N=529) and African (N=756). Briefly, the sample × genotype matrix was scaled to have unit mean and standard deviation along each SNV and subjected to a principal component analysis (PCA). For training the classifier, the genotypes were projected onto the top 6 PCs and used as feature vectors. The classifier is a Multi-layer perceptron with 1 hidden layer, a logistic activation function, L2 regularization term, alpha = 1e-05, size of hidden layer = 6 and a L-BFGS solver. The classifier was implemented using the scikit-learn API in Python. A stratified 10-Fold CV with 80:20 split of the training data was used to tune parameters using a grid search. Cross validation performance on the cohort yielded precision/recall scores as follows: NFE: 0.99/1, AFR: 0.99/1, SAS 0.99/1, EAS: 0.99/1, HIS:0.93/0.97, ME: 0.93/0.77. Samples in this study were subjected to ancestry prediction using the model trained above by projecting their genotype vector to the training PCA model and running the classifier to obtain a given sample’s ancestry probabilities for each of the trained population.

For samples to qualify for a European ancestry analysis, they were required to have a European probability greater than 0.5 and an overall genotyping rate of 0.87 across the 12,840 well-genotyped and ancestry informative markers. Lower genotyping rates were considered as uninformative for ancestry prediction. In the case of low genotyping rate, we considered self-declared ethnicity of ‘White’ as qualifying for the European based analysis.

Furthermore, once the final list was constructed, we applied an additional analysis to control for population stratification by using EIGENSTRAT^32^ to remove samples that were considered as genetic outliers, this ensured that the main cluster of samples was of European origins (see below).

### Cohort construction: CCDS coverage PCA

To account for the variability of the coverage over the CCDS between samples originating from various sequencing kits and platforms, we developed a method to remove samples that are considered outliers due to coverage. This step was performed for samples that passed QC and ancestry prediction filters (if applied), and allowed for maximizing the coding region available for the analysis when harmonizing variant level coverage between cases and controls.

We first randomly selected a set of 1,000 CCDS genes for a coverage test. We next constructed a table where the rows are the samples used for the analysis and the columns are the number of based covered at 10x in each of the 1,000 random genes. Finally, we used the coverage table in a principal-component analysis. Outliers were identified as being further than three standard deviations away from the center of the first four principal components (PCs).

In the Caucasian analysis 3,866 cases and 9,426 controls passed initial QC and ancestry filters and were subjected to the coverage PCA filter. The coverage PCA maintained 3,314 cases and 9,214 controls.

In the diversified population analysis 4,075 cases and 14,494 controls passed initial QC and were subjected to the coverage PCA filter. The coverage PCA maintained 3,468 cases and 13,957 controls.

### Cohort construction: Eigenstrat PCA threshold adjustment

EIGENSTRAT^32^ PCA was used for removing genotypic outlier samples as a final cohort pruning step before running the collapsing analysis. The default threshold for removing outliers is six standard deviations from mean over the top ten PCs. This process, including recalculation of the PCs, was repeated five times.

In the Caucasian analysis 3,208 cases and 8,821 controls passed initial QC, ancestry, coverage PCA and kinship filters and were subjected to the final EIGENSTRAT PCA filter. The EIGENSTRAT PCA maintained the final 3,093 cases and 8,186 controls used for the collapsing analysis, including 383 out of 420 whole genome cases that were mapped by the New-York Genome Center (NYGC).

In the diversified analysis 3,353 cases and 13,373 controls passed initial QC, coverage PCA and kinship filters and were subjected to the final EIGENSTRAT PCA filter. The default EIGENSTRAT PCA threshold removed all 420 NYGC whole genomes. This was the result of the addition to the Caucasian analysis of over 3,000 exomes, which reduced the standard deviation and resulted in the exclusion of NYGC whole genomes in the third PC. As these samples were very high quality and were included in the Caucasian only analysis, we adjusted the threshold of the third PC to seven standard deviations, thus maintaining 402 out of 420 NYGC whole genomes. In total, following stratification phase, we maintained 3,239 cases and 11,808 controls for the collapsing analysis.

### Variant-level quality control

Quality thresholds were set based on previous studies^3,33^. Variants were required to have a quality score of at least 30, quality by depth score of at least 2, genotype quality score of at least 20, read position rank sum of at least -3, mapping quality score of at least 40, mapping quality rank sum greater than -10, and a minimum coverage of at least 10. SNVs had a maximum Fisher’s strand bias of 60, while indels had a maximum of 200. For heterozygous genotypes, the alternative allele ratio was required to be greater than or equal to 25%. Variants were excluded if they were marked by EVS as being failures^34^. Variants were annotated to Ensembl 73 using SnpEff^35^.

### Variant-level statistical analysis

Our primary model was designed to search for non-synonymous coding or canonical splice variants that have a less than 12 cases with a recurring variant in cases and controls (internal MAF) and also a ≤0.1% MAF imposed for each population represented in the ExAC database^16^.

We’ve tested this model in three forms: a standard gene-unit collapsing analysis, a domain-unit analysis and an intolerance-informed gene collapsing analysis. A further gene-based analysis evaluating only rare loss of function (LoF) variants was also performed.

For each of the four models we tested the list of 18,653 CCDS genes. For each gene, we counted the presence of at least one qualifying variant in the gene. A two-tailed Fisher’s exact test (FET) was performed for each gene to compare the rate of cases carrying a qualifying variant compared to the rate in controls. For our study-wide significance threshold, after Bonferroni correction for the number of genes tested across the four non-synonymous models, the study-wide multiplicity-adjusted significance threshold α = (0.05/ [4*18653]) = 6.7×10^−7^. We did not correct for the synonymous (negative control) model.

### OE-ratio intolerance for coding domains

The OE-ratio is calculated using the same approach as the missense tolerance ratio (MTR) that is described by Traynelis et al^19^. This approach uses the observed to expected missense ratio for the 89,522 domain coordinates that are described by Gussow et al.^17^.

For calculating a domain OE-ratio, the following requirements are applied: 1) adequate coverage - at least 50% of the bases within the domain must have at least a 10-fold coverage in the ExAC database, release 0.3^16^. 2) At least five distinct variants (of any annotation) are required to perform a binomial exact test depletion of missense at uncorrected alpha of p<0.05.

There were 67,890 domains that passed the above requirements and were scored for their 0E-ratio. The average size of the remaining unscored domains was usually very short (mean=21bp; median=12) and they accounted for 0.77% of the protein-coding exome. Unscored domains were considered as below the intolerance ratio required for the intolerance-informed analysis (figure 3) to prevent loss of gene level information. 0nce a domain lacking 0E-ratio is implicated in an analysis, its intolerance is examined using the average missense intolerance ratio (MTR)^19^ of the domain in question (http://mtr-viewer.mdhs.unimelb.edu.au).

In the case of *LGALSL*, the last three codons of the coding transcript are a short independent domain that was not mapped to a conserved domain from the CDD. However, this small region is still considered part of the gal-binding domain by other databases^36^. The MTR score for these three codons is below the 30^th^ percentile of intolerance, marking this region at least as intolerant as the implicated galectin binding domain (0E-ratio percentile of 37).

## Acknowledgements

We would like to acknowledge the following groups for contributing ALS samples, sequencing, or clinical data: the New York Genome Center ALS Consortium, including J Kwan, D Sareen, JR Broach, Z Simmons, X Arcila-Londono, EB Lee, VM Van Deerlin, E Fraenkel, LW Ostrow, F Baas, N Zaitlen, JD Berry, A Malaspina, P Fratta, GA Cox, LM Thompson, S Finkbeiner, E Dardiotis, TM Miller, S Chandran, S Pal, E Hornstein, DJ MacGowan, T Heiman-Patterson, MG Hammell, NA Patsopoulos, J Dubnau, A Nath; the ALS Sequencing Consortium, including SH Appel, RH Baloh, RS Bedlack, WK. Chung, S Gibson, JD Glass, TM Miller, SM Pulst, JM. Ravits, E Simpson, WW Xin; the Genomic Translation for ALS Care (GTAC) study, including L Bruijn, S Goutman, Z Simmons, TM Miller, S Chandran, S Pal, G Manousakis, SH Appel, E Simpson, L Wang, RH Baloh, RS Bedlack, D Lacomis, D Sareen, A Sherman, and M Penny.

The collection and sequencing of ALS cases for the New York Genome Center ALS Consortium was funded by the ALS Association and the Tow Foundation. Collection of samples and sequencing for the GTAC study was funded by a partnership of the ALS Association and Biogen Idec. Sequencing of cases for the ALS Sequencing Consortium was funded by Biogen Idec.

We would like to acknowledge the following individuals or groups for the contributions of control samples: The Washington Heights-Inwood Columbia Aging Project (WHICAP). We acknowledge the WHICAP study participants and the WHICAP research and support staff for their contributions to this study; S. Kerns, and H. 0ster. K. Welsh-Bomer, C. Hulette, J. Burke; F. McMahon, N. Akula; D. Valle, J. Hoover-Fong, N. Sobriera; A. Poduri; S. Palmer; R. Buckley; N. Calakos; The Murdock Study Community Registry and Biorepository Pro00011196; National Institute of Allergy and Infectious Diseases Center for HIV/AIDS Vaccine Immunology (CHAVI) (U19-AI067854); CHAVI Funding; R. 0ttman; V. Shashi; E. Holtzman; S. Berkovic, I. Scheffer, B. Grinton; The Epik4K Consortium and Epilepsy Phenome/Genome Project; C. Depondt, S. Sisodiya, G. Cavalleri, N. Delanty; The ALS Sequencing Consortium (see above), The Washington University Neuromuscular Genetics Project; C. Woods, C. Village, K. Schmader, S. McDonald, M. Yanamadala, H. White; G. Nestadt, J. Samuels, Y. Wang; S. Schuman, E. Nading; D. Marchuk; D. Levy; E.

Pras, D. Lancet, Farfel; Y. Jiang; T. Young, K. Whisenhunt; J. Milner; C. Moylan, A. Mae Diehl and M. Abdelmalek; DUHS (Duke University Health System) Nonalcoholic Fatty Liver Disease Research Database and Specimen Repository; M. Winn, R. Gbadegesin; M. Hauser; S. Delaney; A. Need, J. McEvoy; A. Holden, E. Behr; M. Walker; M. Sum; Undiagnosed Diseases Network; National Institute on Aging (R01AG037212, P01AG007232

The collection of control samples and data was funded in part by: Bryan ADRC NIA P30 AG028377; NIH RO1 HD048805; Gilead; D. Murdock; National Institute of Allergy and Infectious Diseases Center for HIV/AIDS Vaccine Immunology (CHAVI) (U19-AI067854); Bill and Melinda Gates Foundation; NINDS Award# RC2NS070344; New York-Presbyterian Hospital; The Columbia University College of Physicians and Surgeons; The Columbia University Medical Center; NIH U54 NS078059; NIH P01 HD080642; The J. Willard and Alice S. Marriott Foundation; The Muscular Dystrophy Association; The Nicholas Nunno Foundation; The JDM Fund for Mitochondrial Research; The Arturo Estopinan TK2 Research Fund; UCB; Epi4K Gene Discovery in Epilepsy study (NINDS U01-NS077303) and The Epilepsy Genome/Phenome Project (EPGP - NINDS U01-NS053998); Biogen; The Ellison Medical Foundation New Scholar award AG-NS-0441-08(to O.C.); B57 SAIC-Fredrick Inc. M11-074; 1R01MH097971 - 01A1; This research was supported in part by funding from The Division of Intramural Research, NIAID, NIH; Funding from the Duke Chancellor’s Discovery Program Research Fund 2014; an American Academy of Child and Adolescent Psychiatry (AACAP) Pilot Research Award; NIMH Grant

RC2MH089915; Endocrine Fellows Foundation Grant; The NIH Clinical and Translational Science Award Program (UL1TR000040); NIH U01HG007672; The Washington Heights Inwood Columbia Aging Project; and The Stanley Institute for Cognitive Genomics at Cold Spring Harbor Laboratory. Data collection and sharing for the WHICAP project (used as controls in this analysis) was supported by The Washington Heights-Inwood Columbia Aging Project (WHICAP, P01AG07232, R01AG037212, RF1AG054023) funded by the National Institute on Aging (NIA) and by The National Center for Advancing Translational Sciences, National Institutes of Health, through Grant Number UL1TR001873. This manuscript has been reviewed by WHICAP investigators for scientific content and consistency of data interpretation with previous WHICAP Study publications.

The content is solely the responsibility of the authors and does not necessarily represent the official views of the National Institutes of Health.

